# Exploring RNA conformational ensembles *in silico*: progress and challenges

**DOI:** 10.64898/2026.02.18.706514

**Authors:** Konstantin Roeder, Guillaume Stirnemann, Louis Meuret, Diego Barquero-Morera, Selene Forget, David J. Wales, Samuela Pasquali

**Affiliations:** Randall Centre for Cell & Molecular Biophysics, King’s College London, Great Maze Pond, SE1 9RL, London, United Kingdom; CPCV Lab, École Normale Supérieure - PSL University, Sorbonnes University, CNRS, 24 rue Lhomond, Paris, France; BFA lab, Paris Cité University, CNRS UMR 8251, INSERM ERL 1133, F-75013 Paris, France; LPENS Lab, École Normale Supérieure - PSL University, Sorbonnes University, CNRS, 24 rue Lhomond, Paris, France; NNF Quantum Computing Programme, Niels Bohr Institute, University of Copenhagen, Blegdamsvej 19-21 Copenhagen, Denmark; Yusuf Hamied Department of Chemistry, University of Cambridge, Lensfield Road, CB2 1EW, Cambridge, United Kingdom

## Abstract

RNA function is intrinsically linked to its structural polymorphism, with molecules exploring the heterogeneous conformational ensembles resulting from complex energy landscapes. These landscapes arise from competing interactions, small energetic separations between microstates, and strong coupling to the environment, posing significant challenges for both experimental characterization and molecular simulation. In this chapter, we review current computational strategies that aim to explore RNA conformational ensembles in silico, with a specific focus on energy landscape–based approaches and atomistic simulations. We discuss key limitations related to sampling efficiency, force-field accuracy, and ensemble analysis, and illustrate their impact through case studies on a self-cleaving ribozyme and an H-type pseudoknot. Finally, we highlight emerging directions, including closer integration with experimental data and the growing role of machine learning, which will probably reinforce the predictive power of in silico RNA energy landscape exploration.

## Introduction

Ribonucleic acids (RNAs) lie at the heart of cellular function, with diverse roles extending far beyond their classical capacity as messengers of genetic information. ^1^ For example, non-coding RNAs (ncRNAs) have emerged as versatile regulators of gene expression, cellular signalling, and metabolic control,^2^ with profound implications for development, physiology, and disease. Their functional repertoire ranges from fine-tuning transcriptional networks to orchestrating complex structural rearrangements that modulate cellular activity, highlighting the necessity of a comprehensive understanding of RNA structural behaviour in biological contexts.^3^ Dysregulation of ncRNA structure and dynamics has also been implicated in disease, from neurological disorders^4^ to cancer,^5^ highlighting the importance of understanding RNA conformational landscapes for therapeutic design.^6,7^

Originally, like for other biomolecules, RNA structure has often been represented by a single, static fold based on the optimal Watson-Crick base pairing. This initial model of RNA structure would allow for relatively straightforward structure prediction,^8^ but neglected the prevalence of non-canonical interactions in RNAs,^9^ which allow for the formation of multiple alternative motifs with comparable energies. Hence, the RNA structural ensembles should rather be considered as polymorphic, sampling a rich variety of configurations across multiple timescales. Pioneering work have indeed highlighted that many functional RNAs are better described by structural polymorphism than by a single dominant conformation. ^10^ The associated energy landscape (EL) exhibit multiple low lying basins stabilizing a series of distinct configurations, potentially with a variety of transition paths connecting them. Importantly, this situation does not mean that the associated energy landscapes exhibit glassy character. Rather, they are multifunneled by stabilising multiple structures, but they still follow the principle of minimal frustration.^11^ As a result, alternative conformations are stable enough (from a kinetic and thermodynamic perspective) to be characterized, allowing for experiments to explore distinct configurations at high resolution. Experiments also reinforce the view that RNA structural ensembles are highly dynamic. NMR and single-molecule approaches have revealed interconversion between states on timescales ranging from picoseconds to seconds.^12,13^ The explicit mapping of alternative states in regulatory RNAs, such as the 7SK 5’ hairpin,^14^ illustrates how energy landscape analysis can uncover functional conformations that would be obscured by single-structure models^15^.

The polymorphism exhibited by RNAs represents a challenge for experimental approaches, especially when it comes to capture short-lived states and slow transitions. Hence computational and theoretical approaches have been important for investigating RNA structural ensembles. Enhanced sampling and molecular dynamics simulations have been applied to both small motifs and larger regulatory RNAs. ^16^ One successful approach to study the dynamics of RNAs lies in exploration of the underlying potential energy surface and calculating emergent thermodynamic and kinetic signatures using statistical mechanics and unimolecular rate theory.^17,18^ Here, the energy landscape is treated as a network of connected stable configurations, revealing links between structure and dynamics. Complementary methods developed by Bussi and Schlick using advanced molecular dynamics and ensemble reconstruction have further refined our understanding of RNA folding pathways and free energy surfaces, illustrating the role of landscape frustration and metastable states in defining RNA polymorphism.^19,20^

These approaches have been applied to biologically relevant systems. For example, an exploration of the energy landscape of HIV-1 TAR reproduced experimentally characterised energy barriers and identified transient conformational states and transition pathways critical for ligand recognition and regulatory interactions. ^21^ Similarly, previous studies explored the 7SK 5’ hairpin stemloop,^14^ the ORF50 transcript from the KSH virus,^22^ and the PK1 pseudoknot,^23^ revealing alternative conformations and folding pathways essential for regulatory function. In viral RNAs, such methods have elucidated structural heterogeneity in SARS-CoV-2 elements. Molecular dynamics and enhanced sampling studies reconstructed diverse ensembles of SARS-CoV-2 RNA,^24^ while Schlick and collaborators focused on the programmed -1 ribosomal frameshifting element (FSE), revealing multiple viable folds, pseudoknots, and interconverting three-way junctions critical for frameshifting efficiency and viral replication.^20,25^ These studies underscore how kinetic pathways connect functional basins, linking structural polymorphism directly to biological outcomes.

Collectively, computational and experimental studies highlight the importance of viewing RNA as a dynamic ensemble rather than a single static structure. A global, energy landscape perspective integrates thermodynamics, kinetics, and structural heterogeneity to explain functional mechanisms, regulatory transitions, and opportunities for targeted therapeutic intervention. This ensemble view is essential, not only for basic biological insight, but also for rational drug design, as demonstrated by studies of viral RNAs, regulatory ncRNAs, and disease-associated RNA motifs.^26,27^ At the same time, significant challenges remain in exploring RNA energy landscapes *in silico*. RNA molecules are highly polymorphic and flexible, with non-canonical motifs and long-range interactions that are difficult to represent accurately with current force fields. Their dynamics span multiple timescales, with significant barriers separating distinct configurations, making it challenging to fully sample all functionally relevant states. Although enhanced sampling methods generally improve results, sampling convergence is non-trivial, with the possibility of missing key states, and connecting simulation results to experimental observables remains challenging.

Here, we discuss these limitations alongside the potential of various approaches to explore energy landscapes, illustrated using two benchmark RNAs that have been extensively investigated. We then provide a perspective on new directions, including strategies to better integrate experimental data into simulations and the opportunities recently opened by artificial intelligence to guide and accelerate the exploration of RNA energy landscapes.

## Challenges of RNA energy landscapes exploration

The computational exploration of RNA structural ensembles is constrained by a combination of methodological challenges, including sampling efficiency, force-field accuracy, and the analysis of high-dimensional structural ensembles.

### Sampling

A widely used and effective description of the existence of multiple biomolecular conformations under given thermodynamic conditions is the free energy landscape (FEL). On the FEL, the relative stability of the different states, and hence their equilibrium occupation probability, is determined by the free energy differences between the corresponding basins, while the kinetics of interconversion between structures are governed by the free energy barriers separating them.

A central and still imperfectly met requirement of molecular simulations is the ability to explore and sample the relevant FEL as thoroughly as possible. This requirement can also be viewed in terms of reproducing a representative sample of all relevant states. Below, we discuss several approaches that can be used for this purpose, without claiming to be exhaustive, and instead focus on the key techniques typically employed as part of our workflows.

Unbiased MD simulations provide direct sampling of the FEL without the introduction of external bias. In principle, they therefore offer the most faithful description of local basin structures and intra-basin fluctuations. This approach is straightforward to implement, as it does not require specific technical choices regarding the definition of collective variables (CVs) and/or the use of enhanced sampling schemes. For RNA systems, such approaches have been important in assessing force-field performance, resolving subtle energetic balances within secondary-structure elements, and characterizing local interaction networks.^16^ However, sampling is typically confined to the vicinity of experimentally determined structures, which can lead to force-field over-stabilization and severely limit the ability of simulations to explore alternative conformational states and the pathways between them. From a landscape perspective, unbiased MD therefore remains intrinsically local: even very long trajectories generally access only a restricted subset of minima connected by relatively low barriers, leaving the global organization of competing basins largely inaccessible.

Biased sampling approaches rely on selecting a set of collective variables (CVs) that effectively represent the slow dynamics of the system. Once chosen, these CVs are used to bias the sampling through methods such as umbrella sampling,^28^ metadynamics and its variants,^29–32^ or techniques that selectively accelerate dynamics,^33,34^ to cite a few representative examples. When the selected CVs correctly capture the reaction coordinates and the remaining degrees of freedom equilibrate rapidly compared to the simulation timescale, these strategies, in principle, enable the investigation of processes irrespective of the magnitude of the associated free energy barriers. However, to explore the conformational landscape of RNAs, identifying approriate CVs is in practice extremely challenging, although an explosion of machine-learning strategies are being proposed to tackle this problem.^35^

Another approach is generalized ensemble sampling, which involves simulating the system under a modified ensemble to artificially accelerate slow transitions. This aim can be accomplished by altering variables such as the temperature^36^ or potential energy.^37^ The thermodynamic properties of the original, unmodified system are typically recovered using replica-based strategies. In these approaches, multiple replicas of the system are subjected to a series of perturbations and periodically exchange configurations, enhancing sampling efficiency and enabling accurate reconstruction of thermodynamic quantities.^36–38^ From a FEL perspective, replica exchange approaches improve coverage across competing basins and reduce kinetic trapping, yielding ensembles that are particularly well suited for thermodynamic analysis and comparative assessment of force fields. However, they do not explicitly resolve transition states or pathway connectivity and remain computationally demanding for large RNA systems. Hamiltonian replica exchange (HREX) is particularly useful and less resource demanding than temperature replica exchange (T-REMD), in particular the solute tempering approach of REST2, which has been used quite extensively for RNA systems.^14,39–44^ While a very recent study suggested faster convergence for T-REMD as compared to HREX when considering the folding/unfolding equilibrium of an 8-mer tetraloop,^45^ T-REMD is computatioannaly challenging to implement for larger RNAs that would require too many replicas.

Ratchet MD occupies an intermediate position between unbiased sampling and more strongly biased approaches. By applying a soft, one-sided bias along a global collective variable, such as the fraction of native contacts, rMD discourages backtracking while preserving much of the underlying local topography. In landscape terms, this bias allows trajectories to preferentially progress toward target basins while retaining realistic intrabasin structure and intermediate minima. For RNA, rMD has proved useful for identifying plausible folding routes and alternative orders of secondary-structure formation, although exploration remains conditioned on proximity to a predefined reference state.^46^

Discrete path sampling (DPS)^47,48^ represents a more explicit approach to explore the energy landscape,^49^ which is described as a coarse-grained network of local minima connected by transition states. The relative energy and densities of states of the minima allows computation of thermodynamic properties, while the transition states enable access to dynamics, considering the set of minima and transition states as a kinetic transition network.^50,51^ Importantly, the framework does not require a reduction in dimensionality, and uses geometric definitions of stationary points, thereby overcoming the connection between sampling time and energy barrier height that classical simulation techniques suffer from. ^52^ This framework is particularly well matched to RNA systems with multiple competing folds, as it enables direct identification of funnels, kinetic traps, and dominant pathways without needing any collective variables. The main limitations for DPS lies in the exponential growth of stationary points with system size, which means that implicit solvent is normally employed for biomolecules. It should be noted, however, that the excellent sampling convergence of the approach allows for a computation of free energy barriers that matching experiment well, limited principally by the shortcomings of the underlying force fields.^21^

### Force Fields

The state of the art in atomistic RNA force fields reflects steady but incremental progress toward balancing structural stability, conformational diversity, and thermodynamic accuracy across heterogeneous RNA motifs. Most contemporary simulations rely on non-polarisable, additive force fields derived from the AMBER and CHARMM families, with AMBER variants (e.g., ff99-based lines with successive *χ*, *α/γ*, *β*, and sugar-pucker refinements) being the most widely used in the RNA community.^16^ These force fields have achieved reasonable stability for canonical A-form helices and improved behaviour for common motifs such as tetraloops and short hairpins, enabling routine microsecond-scale simulations. Nevertheless, even at the current state of the art, RNA force fields remain finely balanced and sensitive: small parameter changes can significantly alter the relative stability of folded, misfolded, and partially unfolded states, underscoring the marginal energetic separation characteristic of RNA energy landscapes. ^53,54^

A central challenge lies in the coupling between backbone conformational energetics and base interactions. RNA force fields must simultaneously describe sugar pucker equilibria, backbone torsional distributions, base stacking, hydrogen-bonding, and electrostatic screening by ions and solvent. While targeted refinements have substantially reduced well-known artifacts (such as spurious backbone transitions or over-stabilization of non-native stacks),^55^ no existing force field consistently reproduces experimental thermodynamics across diverse RNA folds. In particular, noncanonical interactions, base triples, backbone–base hydrogen bonds, and tertiary contacts, remain difficult to capture quantitatively. Errors in these interactions can reshape the energy landscape by overstabilizing specific minima or artificially lowering barriers, with direct consequences for folding pathways and basin connectivity inferred from simulations.^23^ Overall, the current consensus is that modern RNA force fields are sufficient for qualitative landscape exploration, but not yet quantitatively predictive. They can reliably identify dominant basins, characterize structural heterogeneity, and suggest plausible folding routes, especially when combined with enhanced sampling. However, free energy differences between competing states often fall within force-field uncertainty, and conclusions about relative stability or kinetics must be drawn cautiously.

Ion–RNA interactions remain another major limitation at the atomistic level. Divalent cations such as Mg^+2^ play an important role in shaping RNA landscapes, yet are typically treated with simplified non-polarizable models that struggle to capture site-specific binding, competition with monovalent ions, and coupling to conformational rearrangements.^56,57^ As a result, simulations may reproduce global compaction but fail to correctly rank alternative tertiary structures or reproduce experimentally observed ion-dependent free-energy differences. Polarizable force fields and refined ion models offer a promising direction, but their computational cost and limited validation have so far restricted widespread adoption in largescale RNA simulations.^58^ A recent advance by Duboué-Dijon and co-workers introduces a variant of the Amber OLx nucleic acid force field that incorporates implicit electronic polarization via charge scaling of the phosphate backbone (Electronic Continuum Correction, ECC).^59^ This approach improves the description of monovalent and divalent ion interactions with nucleic acids while maintaining the overall conformational properties of RNA, offering a computationally efficient route to more realistic ion-RNA simulations.

Despite significant advances, it is increasingly evident that non-polarizable classical force fields still have fundamental limitations: there is no universal solution to this problem, and performance is often highly system-dependent. Selecting an appropriate force field therefore typically requires careful validation against available experimental data whenever possible. Although polarizable models can offer a more accurate representation of certain critical interactions in these systems, they come with increased computational cost, which can limit sampling efficiency, and these models also demand careful parametrization.

### Ensemble analysis

Beyond issues of sampling efficiency and force-field bias, the rigorous analysis of simulated RNA ensembles remains a significant challenge. RNA structure and function are increasingly understood to arise from ensembles of interconverting conformations, rather than from a single dominant structure, making it essential to develop analysis tools that can robustly extract, summarize, and compare the information encoded in such ensembles. Meaningful characterization therefore requires methods that move beyond individual conformations and purely geometric similarity measures, instead capturing the diversity of interaction patterns and physicochemical environments sampled across the ensemble.

At present, RNA structural analysis tools remain highly fragmented, with most addressing only specific aspects of RNA characterization rather than providing integrated frameworks. Established programs such as VARNA,^60^ MC-Annotate,^61^ and RNApdbee^62^ focus primarily on static structure visualization, annotation, and base-pair classification from single 2D or 3D structures. While highly valuable, these tools are not designed to handle trajectory data or large structural ensembles. More recently, ensemble-oriented tools such as the Python-based Barnaba^63^ have been developed to analyse RNA conformational dynamics from molecular simulations.

Building on the ensemble perspective, we have developed two complementary tools specifically tailored for the analysis and comparison of RNA structural ensembles generated by molecular simulations and energy-landscape–based approaches. The first is the framework of Statistical Molecular Interaction Fields (SMIFs),^64^ which characterizes the three-dimensional space around an RNA molecule in terms of the probability of favourable interactions with possible partners. For RNA, these interactions primarily include electrostatics, hydrogen-bonding (donor and acceptor), and base stacking. By averaging over large numbers of conformations belonging to the same energy basin, SMIFs provide spatially resolved maps of interaction propensities that smooth over thermal fluctuations while retaining chemically meaningful features of the ensemble. This field-based description is particularly well suited to RNA, where conformational heterogeneity, ion-mediated effects, and marginal stability often render single-structure interpretations inadequate.

The second tool is ARNy Plotter,^65^ a web server that integrates widely used RNA structural analysis methods, including Barnaba, into a unified ensemble-oriented framework optimized for trajectory analysis. ARNy Plotter enables the simultaneous characterization of entire structural ensembles, facilitating the comparison of energy basins, intermediate states, and transition trajectories through consistent analysis of base pairing, stacking, and other interaction motifs. When used in conjunction with SMIFs, ARNy Plotter allows population shifts and transient interaction patterns to be directly interpreted in terms of underlying interaction hotspots and landscape features.

Together, SMIFs and ARNy Plotter reflect the state of the art in RNA ensemble analysis by emphasizing statistically robust, interaction-based descriptors, rather than individual conformers. Throughout the remainder of this discussion, both SMIFs and analyses derived from ARNy Plotter will be extensively employed in all reported examples.

## Case studies

Here, we illustrate how different choices of sampling strategies and force fields probe different aspects of RNA energy landscapes using two complex examples: a small self-cleaving ribozyme and a H-pseudoknot.

### Sampling and force-field effects in a minimal self-cleaving ribozyme

Hairpin ribozyme is one of the most thoroughly investigated small self-cleaving ribozymes^66^ first identified in the tobacco ringspot virus, which adopts a catalytic architecture shared with several other ribozymes.^67^ From a simulation standpoint, particular interest has been sparked by reports of multiple pathways associated with markedly different conformations,^68–72^ by the pronounced sensitivity to the choice of force field,^73^ and by the wide range of strategies employed to construct reactive starting configurations from the crystal (methylated) structure.^68–72,74,75^ These features make it a good illustrative system for the present purposes.

#### Comparison of different forcefields

Figure 1 compares the conformational ensembles of the hairpin ribozyme obtained from REST2 simulations on 24 replicas using the OL3 and DES force fields at 300 K, considering the last 200 ns of simulation (out of 400 ns) (see previous work for simulation details^43^). Panel A reports the relative populations of the ten most populated secondary structures. Two clear qualitative differences emerge. First, all top-ten structures in the DES ensemble contain a pseudoknot, whereas the OL3 ensemble also includes highly populated structures without pseudoknots, indicating a weaker stabilization of long-range tertiary contacts. Second, the dominant DES structures display a higher number of canonical base pairs than their OL3 counterparts. In addition, the DES ensemble is dominated by a single secondary structure, while the OL3 ensemble exhibits several alternative 2D structures with comparable populations. Further differences are evident in the conformational landscapes shown in Panel B, where we compute RMSD and percentage of native contacts (Q) with respect to the same structure, chosen as representative from one of the trajectories. Although both force fields sample similar ranges of RMSD and Q values, OL3 displays two well-separated basins, consistent with the coexistence of distinct folded states. In contrast, DES produces a more connected landscape composed of multiple, shallower basins that are mutually accessible.

**Figure 1:**
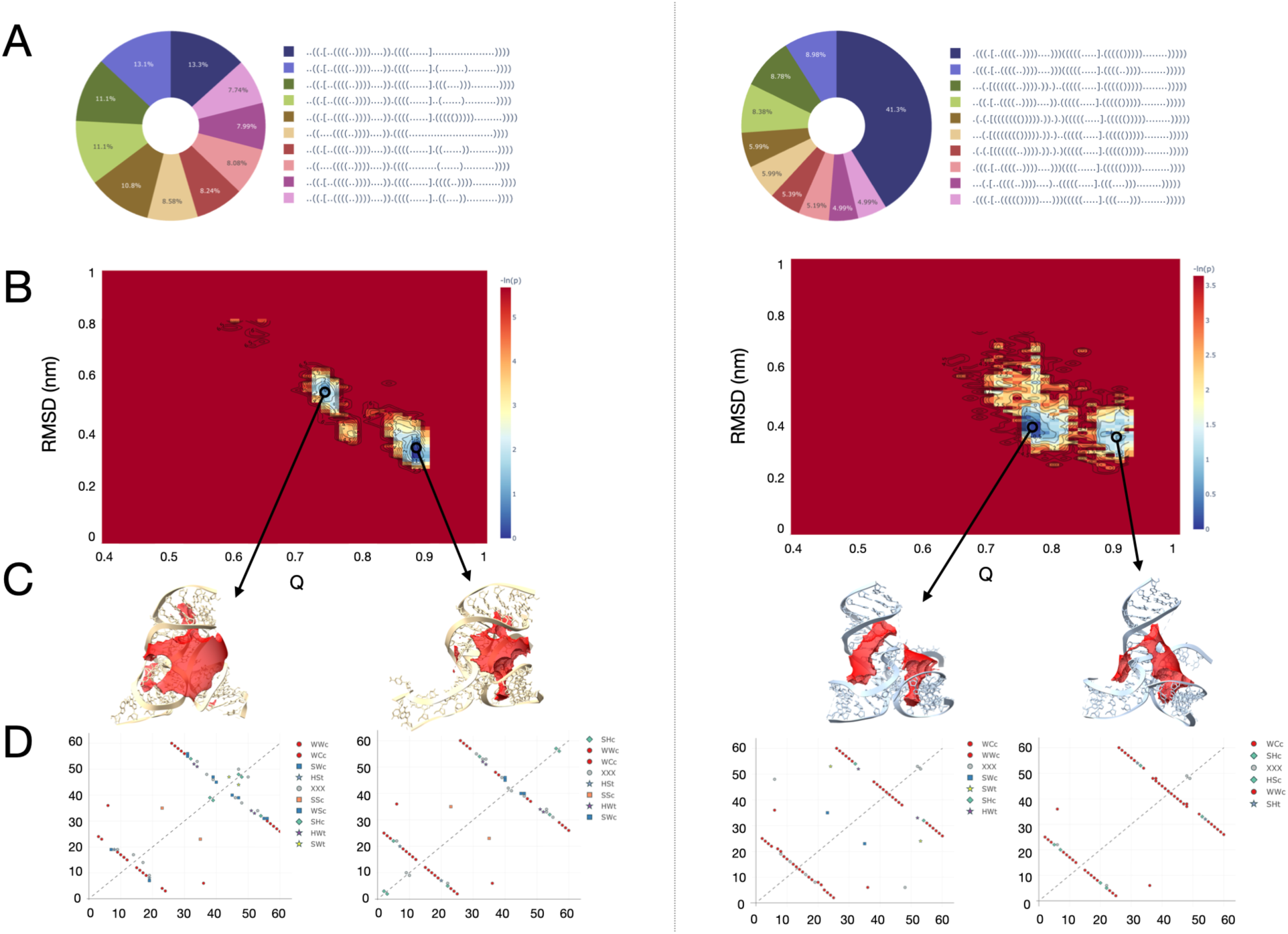
Analysis of REMD simulations of the ribozyme using OL3 (left) and DES force fields (right). A: Relative prevalence of the 10 most populated secondary structures. B: Conformational landscape projected with respect to a chosen reference structure. C: Representative structures of from the most populated basins to which the electrostatic field (APBS) is superimposed, using the same isovalue for all four systems. D: Base pair maps of the four representative structures.

Representative structures extracted from the most populated basins (Panel C) reveal marked differences in the electrostatic environment of the catalytic core, which involves scissible junction bases A5 and G6 and their neighbors G20 and A50. In the OL3 ensemble, the electrostatic field in the core varies substantially between representative structures, whereas it is more conserved in DES. Base-pair maps of the representative structures (Panel D) confirm that OL3 force field promotes non-standard base-pairing interactions, including contacts involving the tertiary structure, whereas DES predominantly stabilizes canonical Watson–Crick base pairs. The base-paring also reveals that the force-field dependence primarily affects interactions within the catalytic core rather than peripheral regions, with notable differences involving A50 and its local base-pairing network. Overall, these results indicate that DES enforces a more structurally and electrostatically homogeneous active-site architecture, while OL3 allows alternative core arrangements, potentially impacting mechanistic interpretations of ribozyme catalysis.

#### Comparison of sampling methods

To assess the effect of enhanced sampling, we performed 24 independent conventional MD simulations at 300 K using the OL3 force field, under the same conditions as the REST2 simulations. Each simulation was initiated from the same reference structure used to analyse REST2 simulations, and extended for 50 ns after equilibration. Therefore, we can compare these to the results of 50 ns of REST2 simulations at a similar computational effort, that is, a fraction of the data used for the above comparison between forcefields. Figure 2 shows the conformational landscape and 2D structures explored by the simulations for both 50 ns of REST2 at 300 K and the 24 MD simulations. We can clearly observe that while REST2 explores the two basins already discussed, even when limiting the time considered, repeated independent conventional MD simulations are restricted to the local free energy minimum defined by their initial structure and do not access alternative states, in contrast to the broader sampling achieved by enhanced methods.

**Figure 2:**
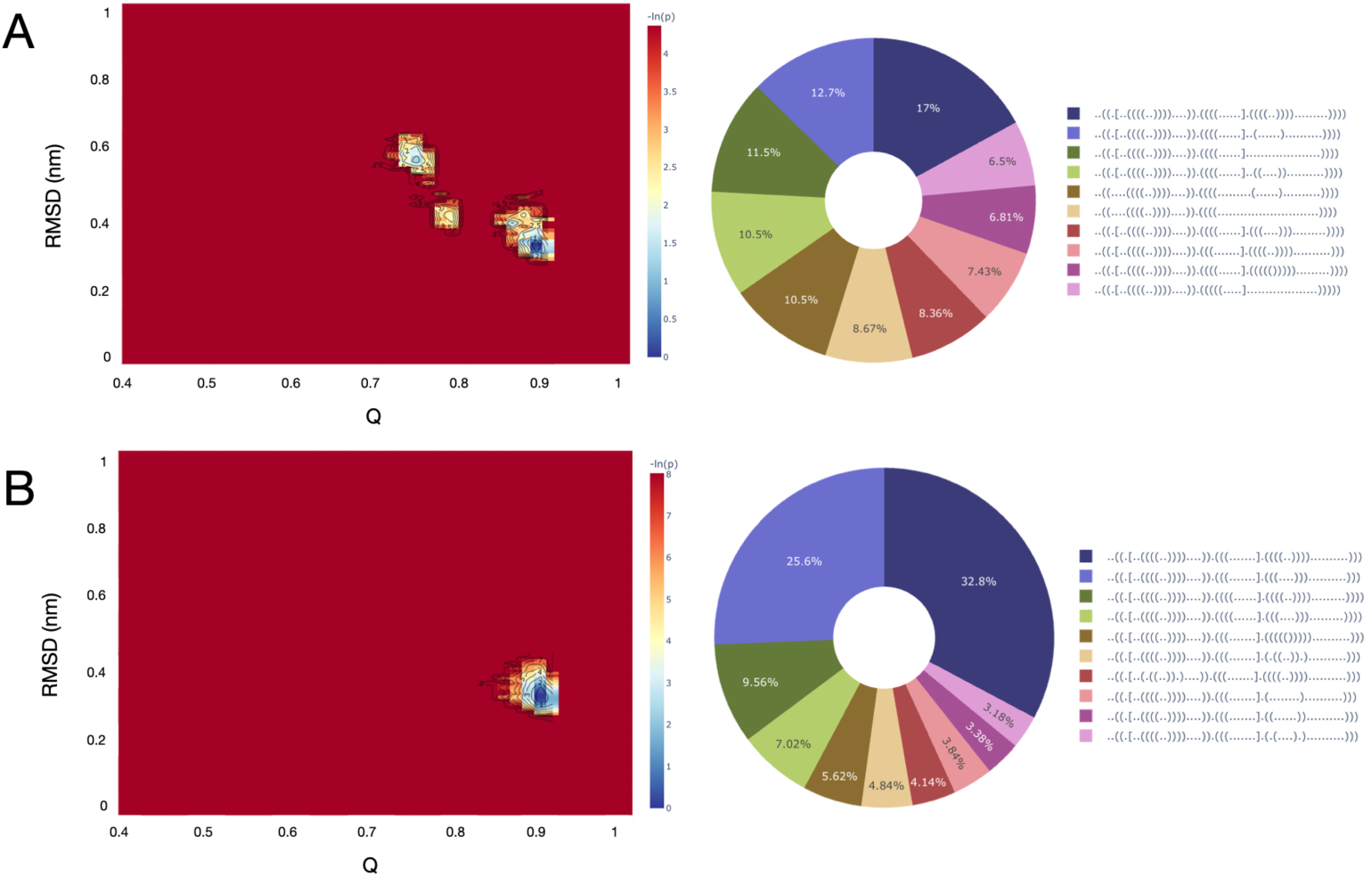
Conformational landscapes and most frequent 2D structures for 50 ns of the 300 K replica of REST2 (A) and for the accumulated 24 MD simulations of 50 ns each (B).

### Energy-landscape exploration of the PK1 pseudoknot

The 22-nt pseudoknot PK1 (PDB ID 2G1W) adopts a compact H-type fold characterized by two interleaved stems and short loops, forming a tight, coaxially stacked architecture stabilized by a mix of canonical Watson–Crick base pairs and non-canonical interactions.^76^ Solution NMR reveals that even this small RNA exhibits a well-defined tertiary core with extensive base stacking and water-mediated hydrogen-bonds that together confer unusually high thermal stability for its size. This structural framework underpins the biological role of PK1 in the Aquifex aeolicus tmRNA, where the pseudoknot promotes ribosomal frameshifting during translation. Its heterogeneous network of canonical and non-canonical contacts makes PK1 an ideal system for probing the energy landscape (EL) of RNA folding and structural dynamics.

#### Comparison of sampling methods

Due to its small size, PK1 has served as an ideal system for exploring RNA energy landscapes and benchmarking both sampling methods and force fields, and we have applied multiple approaches to characterize the conformational space. In 2024, we analysed the EL using discrete path sampling (DPS) across five different force fields,^23^ and separately examined folding pathways with ratchet molecular dynamics (rMD) using the OL3 force field.^46^ DPS simulations with the OL3 force field, widely adopted in RNA studies, revealed a dominant low-energy folded basin corresponding to the native pseudoknot, alongside a set of metastable conformers representing partially folded or misfolded states. These multiple basins reflect the intrinsic frustration of RNA, arising from competing stacking and noncanonical interactions. rMD simulations captured folding heterogeneity by allowing alternative orders of stem formation, yielding ensembles of intermediates and demonstrating the coexistence of parallel folding pathways. In the present study, we extend these analyses with temperature replica-exchange MD (T-REMD). T-REMD simulations were conducted for 200 ns across 26 replicas spanning 278–600 K with the OL3 force field, providing extensive sampling of both folded basins and higher-energy states, which are typically inaccessible to conventional MD on this timescale.

To facilitate direct comparison across methods, we collected three separate datasets: all minima from DPS, productive rMD trajectories, and T-REMD snapshots between 278 K and 360 K. Minima from DPS provide a view of the energetically stable states, without information on transition states or relative populations. Productive rMD trajectories highlight the primary folding routes, though the final folded state is undersampled by design, as the simulation terminates upon reaching this target. T-REMD aggregates configurations across multiple temperatures, offering a qualitative view of the energy landscape by revealing accessible basins and inter-basin regions, but does not yield equilibrium populations at a specific temperature. All three datasets were analysed with ARNy Plotter to examine base pairing, RMSD, fraction of native contacts (Q), and the overall conformational organization of PK1. Figure 3 presents low-dimensional projections of PK1’s energy landscape using DPS, rMD, and T-REMD, mapped onto Q and RMSD. All projections reveal a rugged landscape with multiple competing basins of similar depth, reflecting the shallow energetic separations and intrinsic frustration characteristic of RNA folding.

**Figure 3:**
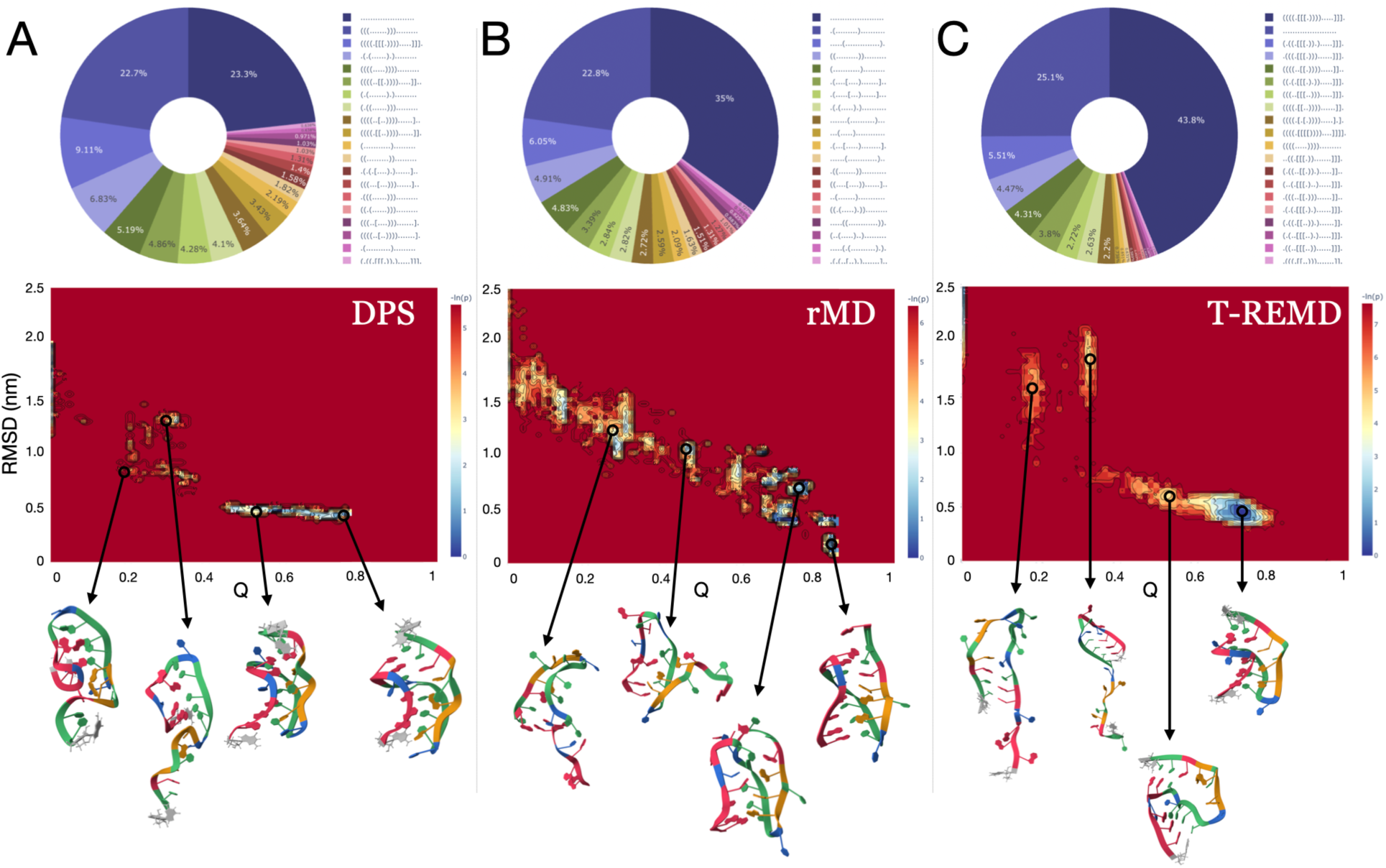
Comparison of PK1 conformational ensembles obtained with the three different methods DPS, rMD, T-REMD. Top: 20 most populated secondary structures and their relative presence. Middle: Conformational landscape projected onto RMSD and Q-values of the experimental structure. Bottom: 3D structures representatives of the most populated states explored by each method.

In the DPS projection (Figure 3A), the conformational landscape is resolved into a set of discrete minima corresponding to stable conformations. The sparse distribution of points in the Q–RMSD plane reflects the identification of distinct basins rather than a continuous sampling of configurations. This representation highlights the presence of metastable states. The most populated 2D structural motifs are distributed across the identified minima and include both fully folded and partially folded states. The native pseudoknot and the single-stem conformation are the two most frequent structures (excluding the fully unfolded state), although several other partially folded conformations are also sampled. Representative structures exhibit substantial structural diversity, including partially formed pseudoknots and misfolded topologies.

The rMD projection (Figure 3B) displays a more continuous distribution along increasing Q-values, emphasizing dominant folding routes and the connectivity between intermediate states. The most populated two-dimensional structural motifs are partially unfolded, consistent with the fact that rMD does not sample the native state. These motifs include both pseudoknotted architectures and simple stem structures with a comparable number of native contacts, suggesting that these arrangements may belong to alternative folding pathways. Representative structures reveal a progressive assembly of the pseudoknot along these routes. T-REMD projections (Figure 3C) exhibit well-defined population maxima associated with metastable basins and reveal regions connecting these basins that are explored through barrier-crossing events. The corresponding two-dimensional structural motifs are largely dominated by the folded state, which is used as the starting configuration for the simulations. The second most populated state is the fully unfolded conformation, with intermediate states sampled much less frequently.

Together, these methods provide complementary views of the PK1 energy landscape. DPS maps the global organization of stable minima and folding funnels; rMD highlights directed folding pathways and intermediate connectivity; and T-REMD captures accessible basins and broader exploration of the landscape. Integrating these approaches gives a comprehensive perspective of the PK1 rugged energy landscape, revealing folding intermediates, metastable states, and the dominant structural motifs connecting them.

#### DPS and experimental data

Our recent systematic comparison of five widely used AMBER RNA force fields using explicit energy landscape explorations provides a detailed view of how current atomistic potentials shape RNA conformational ensembles beyond the native basin.^23^ Using PK1 pseudoknot as a test case, the results show that all five force fields identify the experimentally observed pseudoknot fold as a low-energy structure, even in the absence of explicit ions. The native basin is therefore qualitatively reproduced across models, with broadly similar secondary structure features. Nevertheless, significant differences emerge in the detailed interaction networks, the extent of stacking versus non-canonical base pairing, and, most importantly, the organization of higher-energy regions of the landscape.

At the level of energy landscape topology, the force fields diverge markedly. Some predict clear partially folded intermediates stabilized by individual helical elements, while others strongly destabilize such states or instead favour alternative, non-native folds. These differences lead to qualitatively distinct folding scenarios, ranging from stepwise assembly to highly cooperative folding, with barrier heights between basins differing by tens of kcal/mol between force fields. Such variations are not apparent from local MD simulations near the native state, but become evident only through global landscape exploration, highlighting the sensitivity of RNA folding mechanisms to force-field parametrization.

For this system, the relevance of these differences can be tested directly through thermodynamic observables derived from the landscapes, in particular heat-capacity (C***_v_***) or melting curves (Figure 4). We focus in particular at the qualitative shape of the melting transition, which is a feature directly linked to the energy landscape topology. Two independent experimental studies report a single cooperative melting transition, indicating the absence of stable, long-lived metastable intermediates. The first study was carried out together with the experimental determination of the NMR structure of the system (data not published at the time). The melting transition was monitored with UV spectroscopy at two separate Mg^+2^ concentrations, measuring absorbance. The second was performed by our collaborators, using differential scanning fluorimetry. When C***_v_*** profiles are computed from the energy landscapes from our DPS simulations,^1^ only one force field reproduces this experimental signature. While several of them produce multiple C***_v_*** peaks, reflecting the presence of distinct and thermodynamically stable partially folded states, OL3 uniquely predicts a single dominant peak, consistent with a two-state cooperative transition. Because of the approximations in extracting melting curves from DPS simulations, the melting temperature is an estimate, but the presence of a single or multiple peaks in the curve should be reliable.

**Figure 4:**
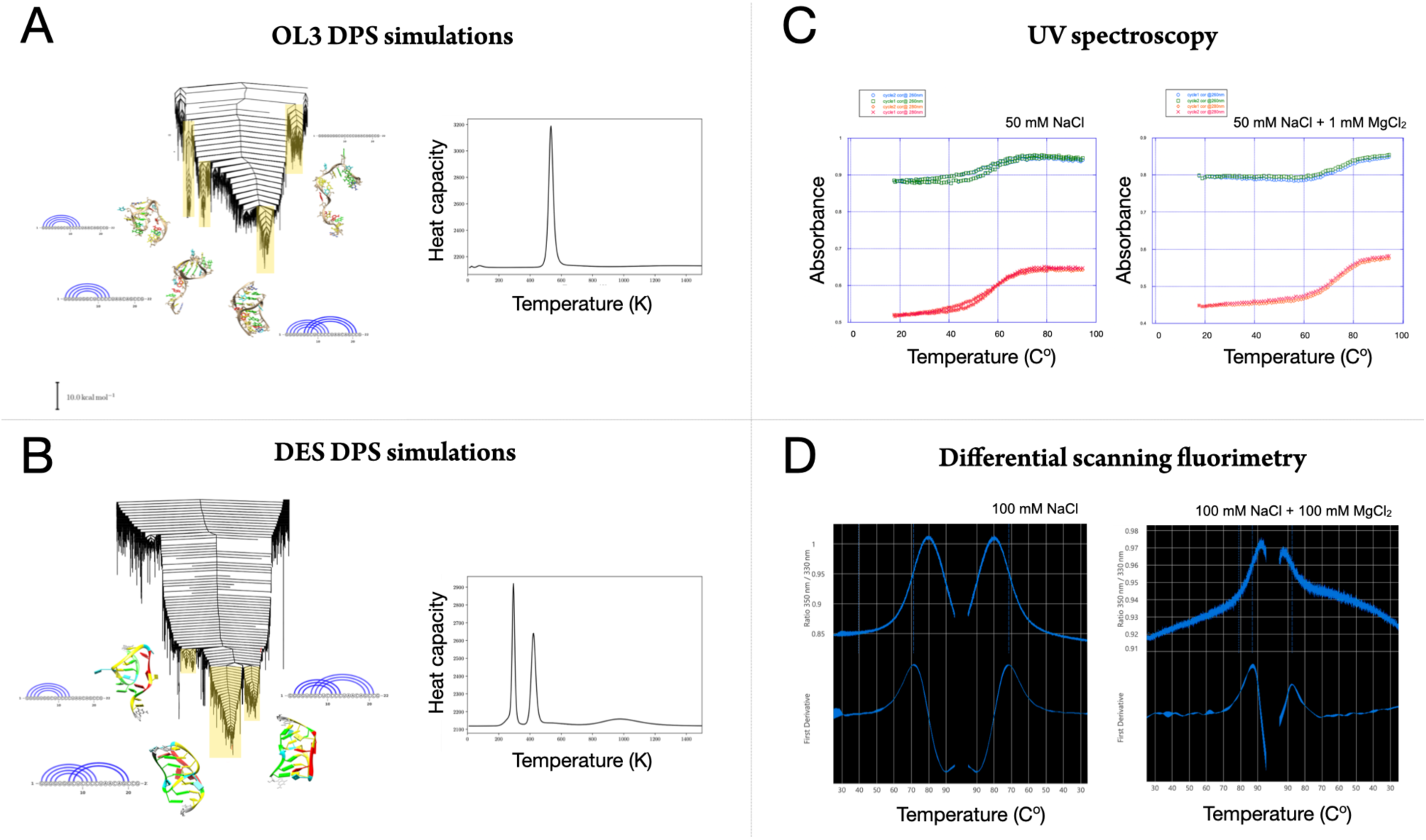
Left: Disconnectivity graph and C***_v_*** curve of PK1 obtained by DPS simulations with the OL3 (A) and DES force fields (B). Right: 2006 experimental data showing the melting transition for Pk1 by UV absorbance (C) and new curves obtained from differential scanning fluorimetry (D) to monitor the melting transition.

Taken together, these results demonstrate that current RNA force fields can agree on native structure yet differ profoundly in their predictions of ensemble thermodynamics and folding mechanisms. The fact that, for the system in question, the OL3 force field is consistent with the experimentally observed single melting transition is reassuring, particularly given its widespread use within the community. At the same time, the failure of other force fields to reproduce this basic thermodynamic feature highlights the need to evaluate RNA force fields against global landscape properties and experimentally accessible ensemble observables, rather than relying solely on native-state stability or short-timescale behaviour.

#### Effects of different charge parametrizations

To assess the impact of implicit electronic polarization on RNA conformational dynamics and ion-mediated interactions, we compared two atomistic MD simulations performed with the standard Amber OL3 force field and with its ECC-modified variant. We performed two molecular dynamics simulations of of PK1 of 200 ns each at 310 K and in solution with the same ionic conditions (150 mM K^+^ and Cl*^−^* and 5 mM Mg^+2^), one using the standard OL3 force field and one using ECC-modified force field. Structural ensembles and interaction fingerprints were analysed to evaluate both global conformational stability and the local interaction landscape.

Both simulations remained structurally stable over the full trajectory, with no evidence of large-scale unfolding (Figure 5). However, clear differences emerge in the temporal evolution of the RMSD. In the OL3 simulation (Figure 5A), the RMSD exhibits pronounced fluctuations and a detectable shift between distinct conformational regimes, suggesting the presence of multiple distinct states sampled over the course of the trajectory. In contrast, the ECC-corrected simulation (Figure 5B) displays a narrower RMSD distribution with reduced long-timescale drift, indicative of a more homogeneous ensemble and enhanced conformational stabilization. This behaviour is consistent with a smoother coupling between RNA structure and the ionic environment when polarization effects are implicitly included. The base-pairing distribution also differs between the two simulations (Figure 5A and 5B, right panel), suggesting that improved treatment of ion–RNA interactions can shift the balance between alternative secondary and tertiary structures.

**Figure 5:**
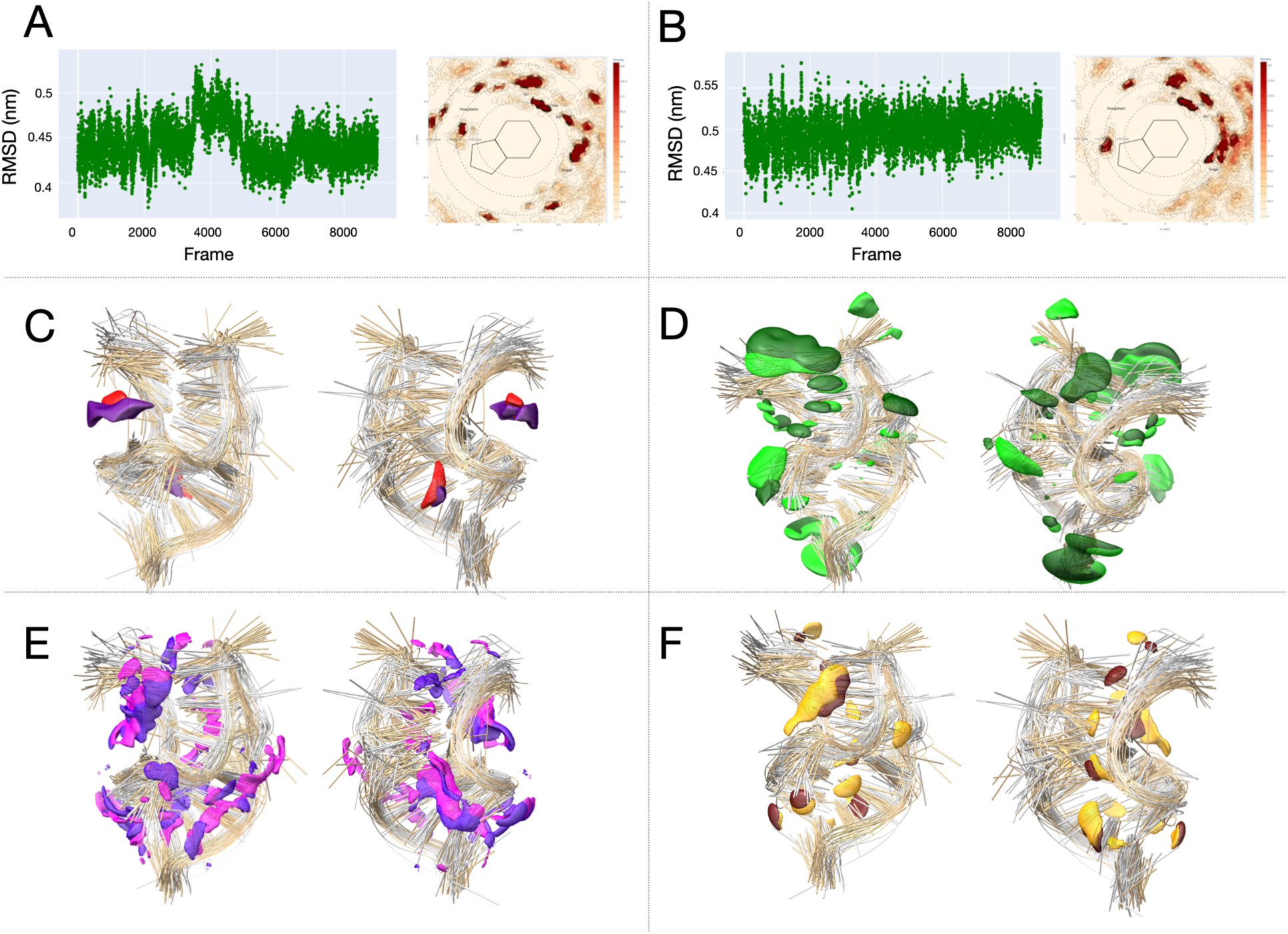
A, B: RMSD (left) and base-pair spatial distribution (right) for OL3 MD (A) ECC (B) simulations. C,D,E,F: Superposition of the structures explored by simple MD using OL3 (grey) and with ECC (brown). Average SMIFs are represented together with the structures and are shown from two viewpoints corresponding to a rotation of 180 degrees around the vertical axis: negative electrostatic (C), Stacking (D), Hydrogen bond acceptors (E), Hydrogen bond donors (F). OL3 SMIFs represented in lighter shades and ECC in darker shades. For each kind of field, the same isovalues are used to visualize SMIFs.

A more detailed picture emerges from the analysis of Statistical Molecular Interaction Fields averaged over the whole trajectory (average SMIFs), which report the spatial distribution of interaction hotspots around the RNA. While both force fields explore broadly similar regions of conformational space, the nature and localization of interaction patterns differ. Negative electrostatic interaction regions (Figure 5C) are more spatially focused in the ECC simulation, whereas the standard OL3 force field yields more diffuse and fragmented electrostatic hotspots. This suggests that charge scaling of the phosphate backbone reduces overly strong, non-specific ion–phosphate contacts and favours more localized electrostatic stabilization. Stacking interaction fields (Figure 5D) are also redistributed upon ECC correction, with clearer and more persistent stacking regions observed relative to OL3. This result indicates that improved ion screening indirectly modulates base–base interactions by stabilizing specific RNA geometries. Hydrogen-bond acceptor (Figure 5E) and donor (Figure 5F) SMIFs show a reorganization toward fewer interaction sites in the ECC simulation. In contrast, OL3 produces broader interaction volumes, consistent with excessive ion residence and overpopulation of hydrogen-bonding configurations. The ECC correction therefore appears to sharpen the specificity of ion-mediated hydrogen-bonding, reducing artificial stabilization of transient contacts.

Taken together, these analyses indicate that incorporating implicit polarization via ECC does not dramatically alter the global fold of the RNA, but significantly reshapes the underlying interaction landscape. The ECC-modified force field promotes more localized, structured, and persistent interaction motifs while suppressing diffuse, non-specific ion binding. This suppression results in a smoother conformational ensemble with reduced structural drift and improved discrimination between competing interaction patterns. These results support the hypothesis that ECC provides a computationally efficient means to enhance the physical realism of ion–RNA coupling without compromising the intrinsic conformational properties of RNA.

## New strategies

Two main strategies are emerging alongside steady improvements of existing simulation methods. First, experimental approaches are more and more integrated directly into computational work. Second, the advance of data-driven methods, i.e. machine-learning (ML) approaches, has started to make its mark on RNA structure prediction.

### Integration of simulation and experiments

The recognition that RNA structure is highly dynamic and relies heavily on non-canonical interactions is also reflected in a shift of how structure prediction in 2D and 3D is conceived and combined with experiment.

RNA secondary structure (‘2D predictions’), i.e. the determination of all canonical Watson Crick base pairs formed, is fundamental to our understanding of RNA structure. The large number of non-canonical interactions builds on this framework to form complex, three-dimensional structures. Computational work has shown that such non-canonical interactions can be relevant to determine the correct binding motif,^14^ or stabilise intermediates, lowering the free energy barriers for structural transitions,^21^ amongst many other key roles. The most suitable approach for RNA structural ensemble studies is therefore to integrate 2D and 3D predictions together with experiment. While the techniques for each of these three areas are distinct and require significant knowledge and expertise, they individually are not currently sufficient for rapid, high resolution of the RNA structural ensembles. Integrated studies are therefore the new frontier in RNA structural ensemble studies, drawing on the strengths of different approaches,^78^ and an example was recently published by Thiel et al.^79^ A combination of data and tools from experiment (SAXS data), secondary structure prediction and 3D modelling was used to map RNA structural ensembles that are sampled globally and locally, and match experimental data.

### Machine-learning potentials

As already discussed, the development of accurate force fields for RNA simulations remains highly challenging. These challenges stem both from the adequacy of the chosen functional form and level of theory (e.g., fully atomistic versus coarse-grained, polarizable versus non-polarizable models) with respect to the phenomena under investigation, and from the difficulty of determining parametrizations that faithfully reproduce ab initio (i.e., quantum-mechanical) reference data and/or experimental results.

Machine-learned interatomic potentials (MLIPs) are now being used in molecular simulations to fit molecular interactions derived from quantum calculations, at a fraction of the associated computational cost.^80–82^ While this technology has already influenced materials science^83,84^ and shows great promise for the study of complex chemical reactivity,^85,86^ applications to biomolecular simulations are still limited. Nevertheless, very promising early studies have demonstrated the feasibility of applying MLIPs to simulations of solvated biomolecules^87,88^ although significant challenges remain, including the incorporation of explicit electrostatics, improvements in computational performance, and the construction of high-quality training datasets for reference, so-called ‘foundation models’, for biomolecular simulations.

In parallel, another active line of research explores the use of machine learning to improve the calibration of coarse-grained force-field parameters. ^89^

### Machine-learning for RNA structure prediction

The use of ML tools to predict RNA structural ensembles comes in many different forms, and there is, at the time of writing, no one tool that enables a prediction of the full polymorphic ensemble for RNAs. There are multiple different approaches that have been developed, broadly falling into three (sometimes complementary) categories: a) the prediction of RNA structure similar to the prediction of protein structures, for example by AlphaFold or RosettaFold, b) the prediction of the ‘full’ ensemble by producing a representative sample of structures, and c) the use of ML tools within existing enhanced sampling workflows, reducing computational cost by faster sampling convergence.

There are a number of challenges to using ML tools for RNA structure prediction. The first question relates to the most fundamental point. Is the prediction of a single structure useful in the context of simulations of RNA structural ensembles? In our opinion, it is a useful tool that may provide reasonable starting structures for further explorations. However, such a single structure is not sufficient to solve RNA structure prediction. Due to the polymorphism exhibited by RNAs, more than one structure is required to describe the structural ensemble, as detailed above. A second challenge lies in the importance of environmental conditions, such as the presence of ions, on RNA dynamics and structure. Predictions should include this sensitivity. A final challenge is the paucity of high quality data for RNA structures. This sparsity reduces the ability to train ML models, and, due to the likely underrepresentation of shorter lived structures, may bias towards configurations with canonical base pairing.

Nonetheless, several ML tools are able to produce good predictions for large RNAs. Some of these tools are not specific to RNA, with AlphaFold3 and Boltz2 as prominent examples. While some benchmarks exhibit reasonable success in predicting RNA structures, the most recent CASP competition found that the top performance still derives from human predictors.^90^ The CASp competition for nucleic acid structure also highlighted already discussed problems with predicting non-canonical interactions, and the impacts of the environment. A key conclusion of the authors is the need for “novel approaches unique to nucleic acids”.^90^

Attempts to predict complete biomolecular ensembles remain relatively limited, and examples involving RNA are particularly scarce. Nevertheless, we suggest that such approaches are especially well suited to polymorphic RNA structural ensembles. Although not all machine-learning methodologies are built around this principle, a conceptually appealing strategy is to exploit modern generative models to map a difficult-to-sample conformational space onto a more tractable latent distribution. This can be achieved using techniques such as normalizing flows, diffusion models, or related approaches.

Despite their expressive power, these methods are fundamentally constrained by the quality of the training data. Consequently, beyond the fine-tuning of model architectures, substantial effort has been devoted to the design of strategies for generating appropriate training datasets. While a comprehensive review of this rapidly growing body of work is beyond the scope of this discussion, we highlight below in an oversimplistic way two representative approaches that address these challenges.

The first tool in this category is BioEmu^91^ (currently available only for proteins). The generative model is initially pre-trained on the AlphaFold Structural Database. An intermediate model is then trained using a large collection of MD trajectories, followed by fine-tuning on experimental stability data to produce the final model. During inference for a specific target system, the model generates a conformational ensemble but does not require additional simulations per se.

Another approach we highlight here is the very recent RNAnneal,^92^ which specifically targets RNA. It operates in several stages: candidate generation using conventional 2D and 3D structure prediction methods, MD simulations of the resulting candidates, and the development of an RNAnneal score based on a physics-inspired generative model. This model, essentially a diffusion model trained across a range of thermodynamic condition that captures the log-likelihood of a conformation, is used to rank candidate structures and ultimately to classify conformations, distinguishing plausible experimentally resolved conformations (ERCs) from decoys. As opposed to BioEmu, a different model is trained for each specific target system.

Finally, ML can be leveraged in a wide range of contexts to accelerate the convergence of MD simulations. For instance, the application of ML to improve enhanced sampling strategies in MD is an area of intense and rapidly expanding research, and a comprehensive review is beyond the scope of this work. We therefore refer the reader to several excellent recent reviews on the subject.^93–95^ Existing approaches span the use of ML to identify optimal low-dimensional representations, the design and optimization of biasing strategies, and the complementing (or in some cases replacement) of traditional sampling methods with generative approaches. It is our opinion, that generative approaches ought to be grounded in physics to be relevant.^96,97^ For example, a tight interplay and iterative feedback between generative samplers and physics-based, exploration^98^ or the development of biased generative samplers,^99^ holds great promise, even though substantial progress remains to be made.

## Conclusions

RNA molecules operate on complex yet highly organized energy landscapes, with relatively high barriers between well-defined functional states that encode functional polymorphism, kinetic control, and response to cellular context. As we have discussed, neither static structural models nor short-timescale simulations are sufficient to capture this complexity. Instead, a landscape-based perspective, integrating thermodynamics, kinetics, and ensemble heterogeneity, is essential to understand how RNA structure is related to function. Through two examples, we have illustrated how different sampling strategies and force fields reveal complementary aspects of RNA landscapes, from dominant basins and folding funnels to metastable intermediates and transition pathways. At the same time, these systems highlight that present-day simulations remain limited by incomplete sampling, force-field inaccuracies, and challenges in ensemble analysis, such that quantitative predictions must still be interpreted with caution.

Future progress in RNA energy landscape exploration will depend on tighter integration across methodologies. Advances in enhanced sampling, landscape-based techniques, and ensemble analysis tools are steadily improving our ability to characterize RNA polymorphism in silico, but their full potential will only be realized through systematic validation against experimental ensemble observables. In parallel, emerging strategies, ranging from improved treatments of ion-RNA interactions to machine-learning approaches for force-field development, structure prediction, and sampling acceleration, offer promising avenues to overcome current bottlenecks. Rather than replacing physics-based models, these data-driven methods are most powerful when embedded within a physically grounded framework. Ultimately, a synergistic combination of simulations, experiments, and machine learning will be required to achieve predictive, ensembles descriptions of RNA structure and dynamics, enabling a deeper mechanistic understanding of RNA function and more reliable avenues for RNA-targeted therapeutic design.

## Acknowledgement

The authors thank Dr S. Nonin-Lecomte for retrieving and sharing with us the UV absorbance data of PK1 as well as Dr Filippo Prischi for performing the newest measurement of PK1 melting curves. SP thanks the French National Research Agency MERLIN ANR-22-CE45-0032 for financial support.

## Supporting Information Available

All simulation trajectories and SMIFs are available at DOI:10.5281/zenodo.18607810 (Full PK1 force field comparisons are available at DOI:10.5281/zenodo.10590336).

## Footnotes

1 The C*_v_* curves are computed by applying a harmonic approximation to obtain vibrational contributions to the partition function. ^77^

